## Supplementary figures and images for "Ultraslow serotonin oscillations in the hippocampus delineate substates across NREM and waking"

### Supplementary Figure 1

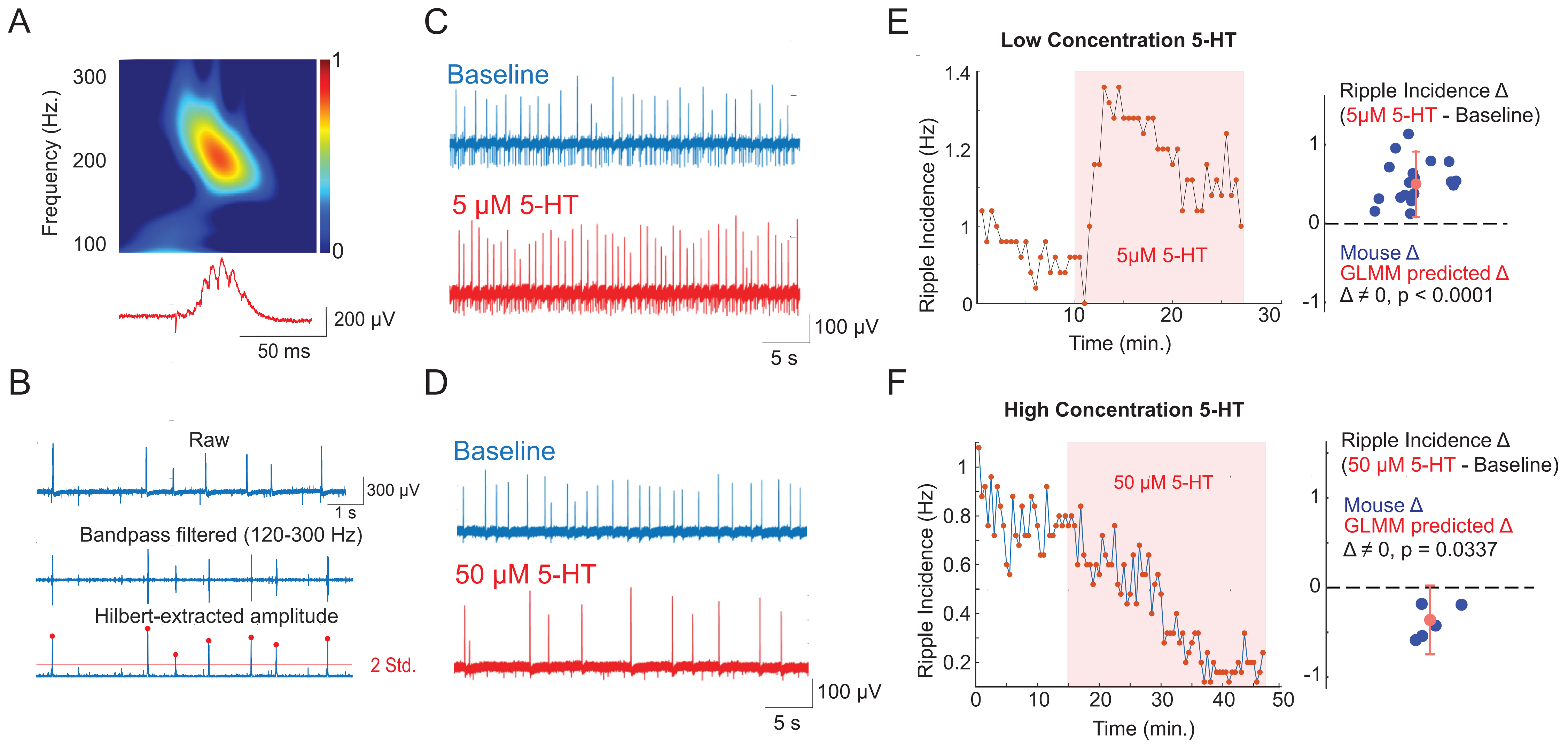

### Supplementary Figure 2

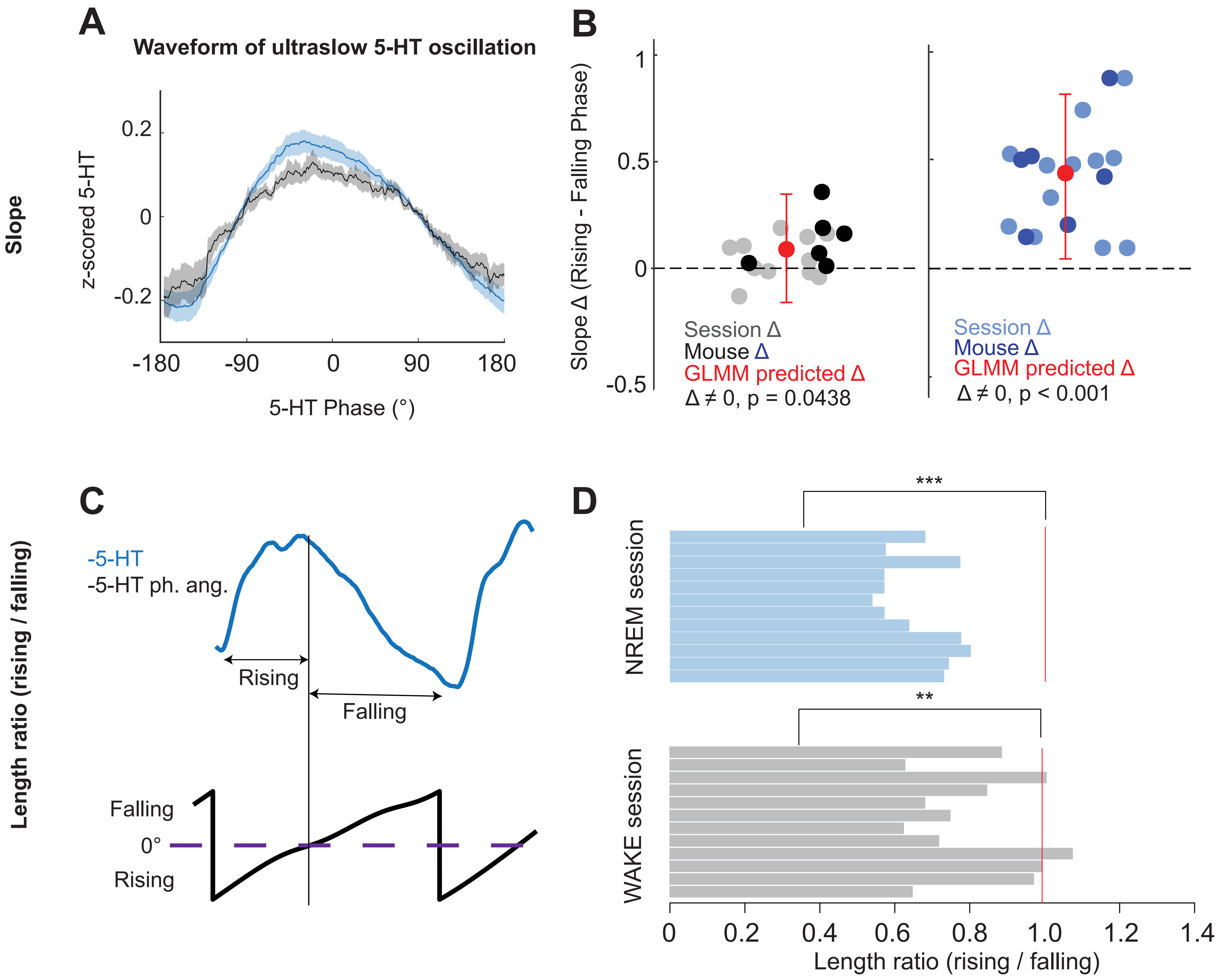

### Supplementary Figure 3

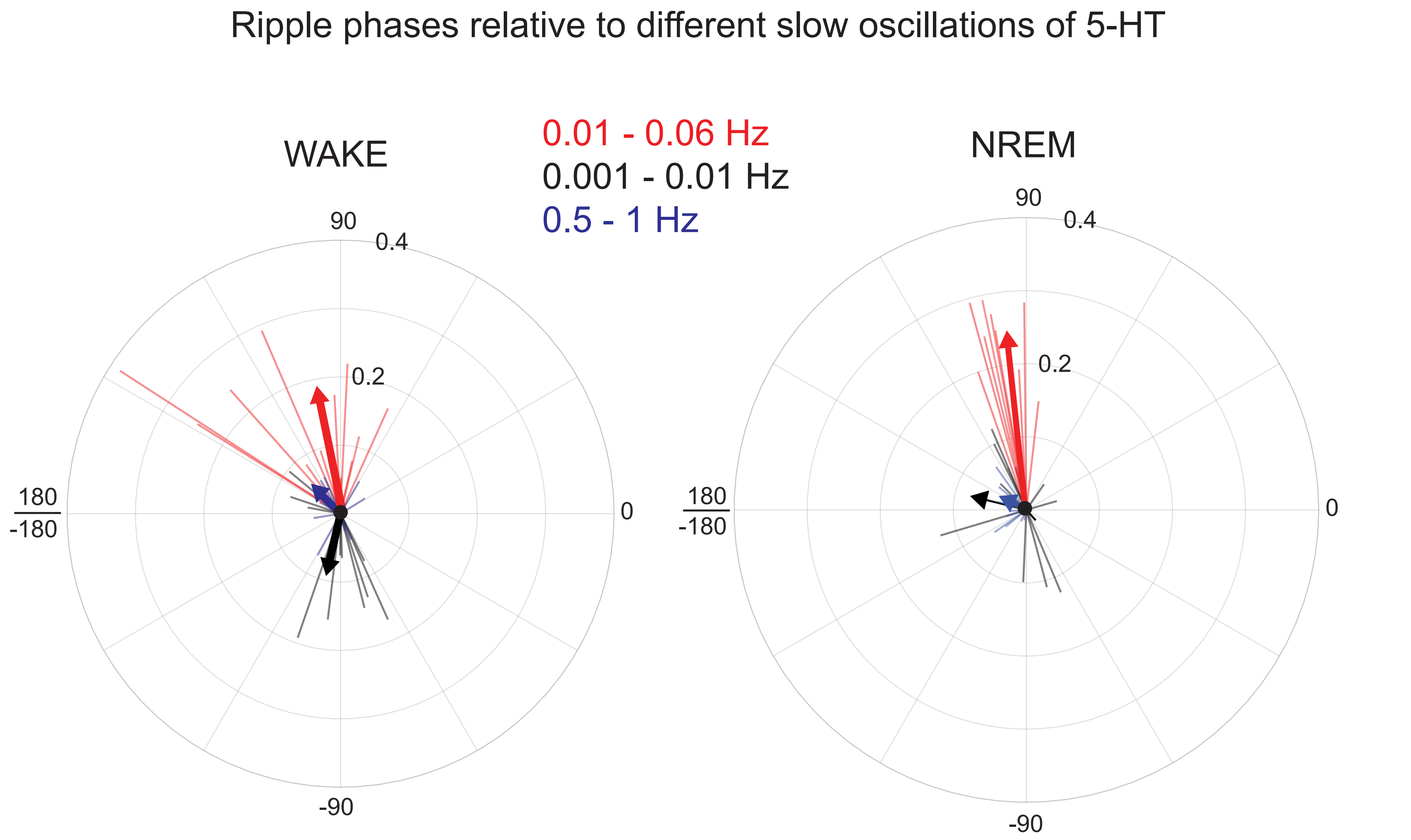

### Supplementary Figure 5

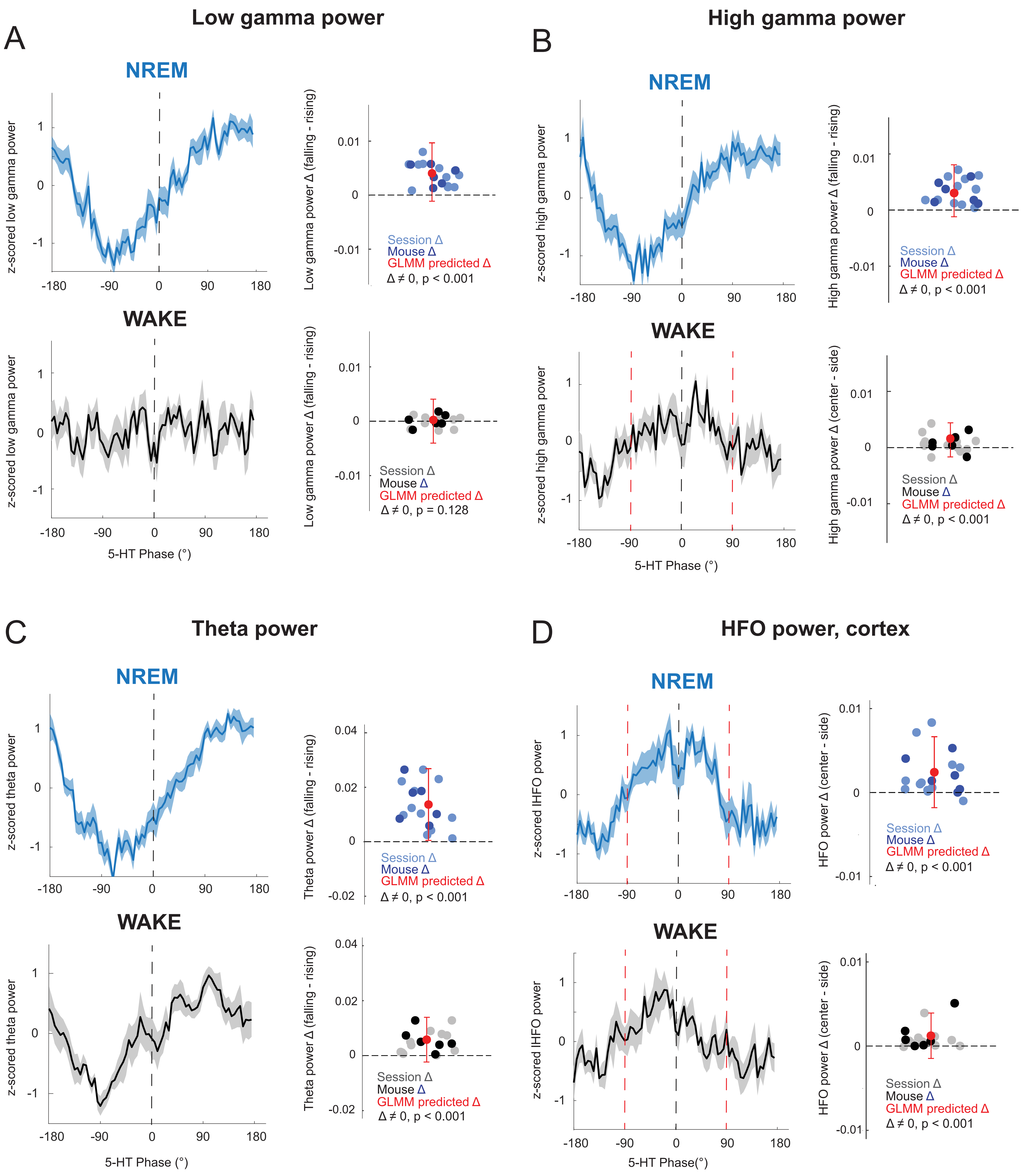

### Supplementary Figure 6

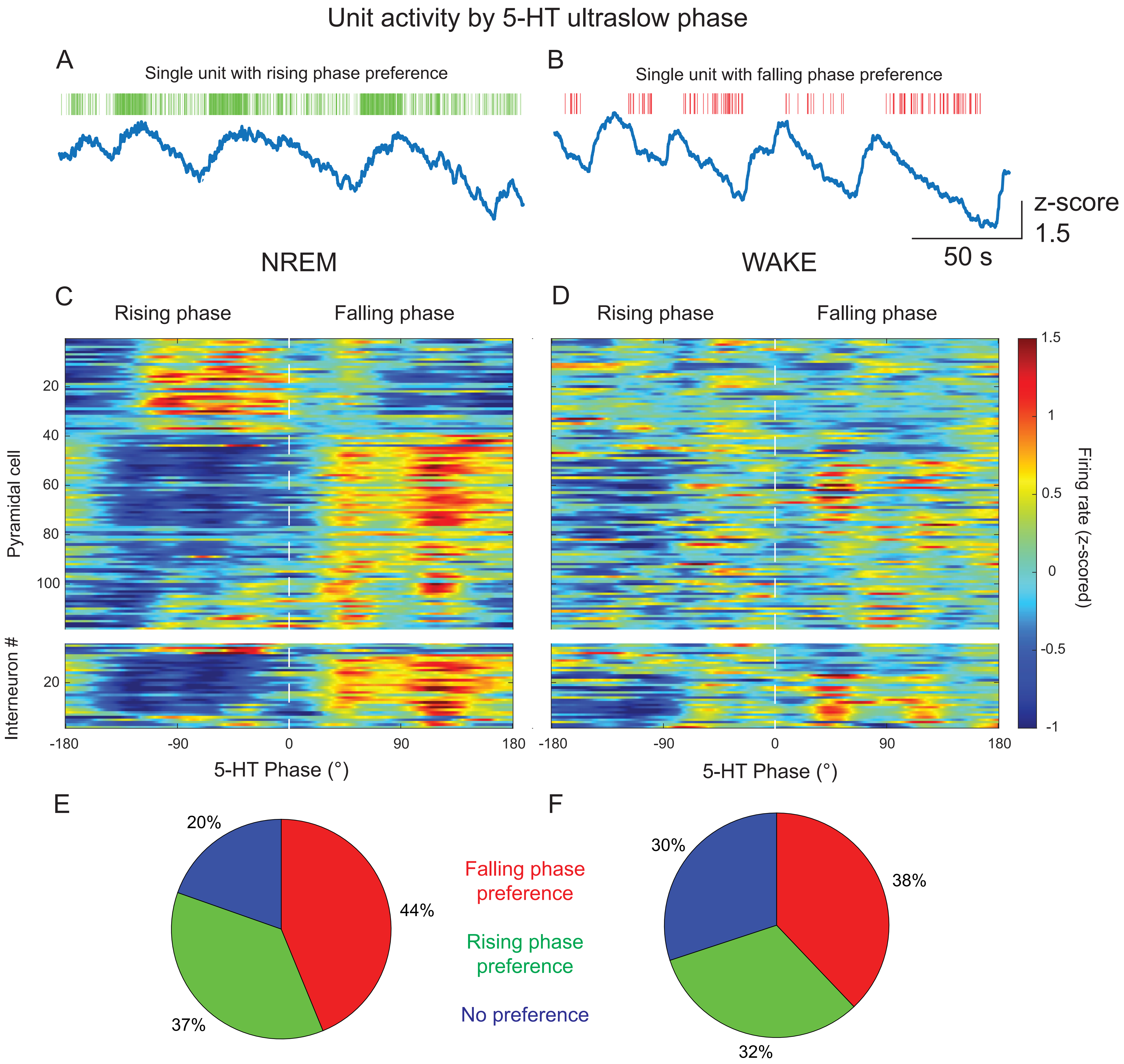

### Supplementary Figure 7

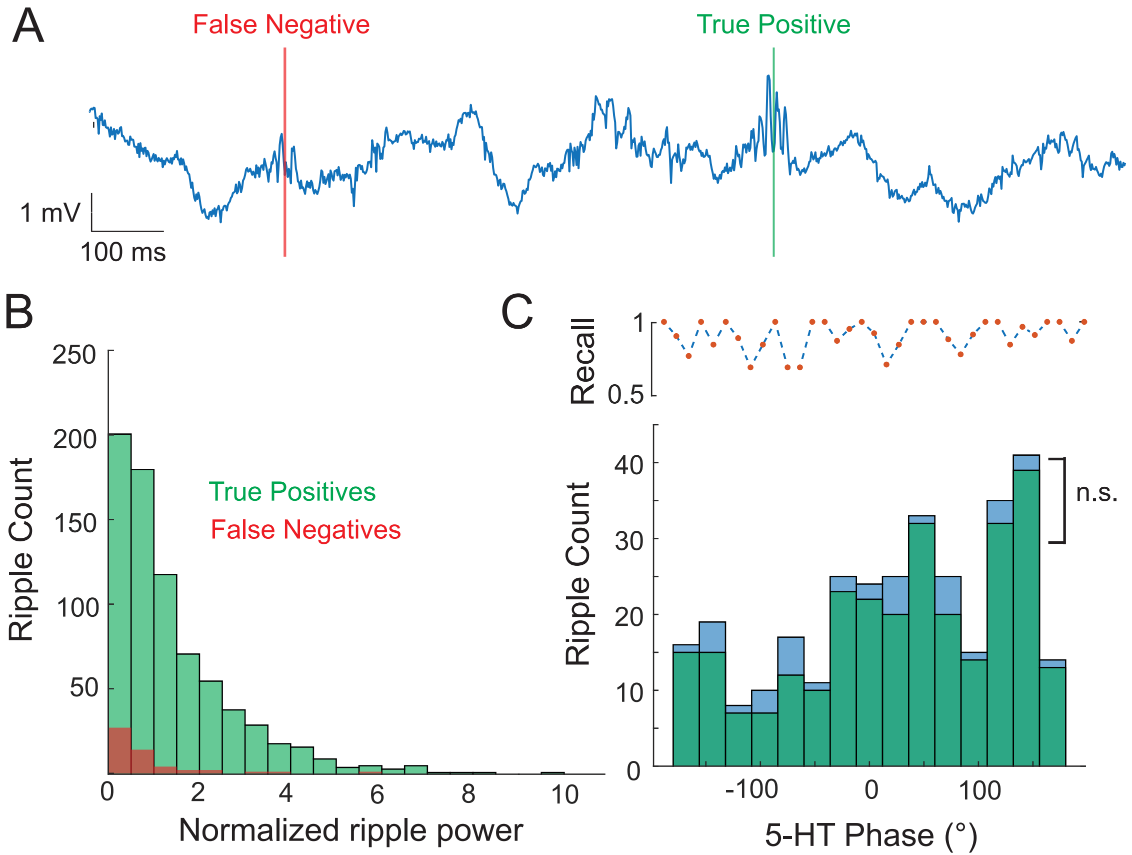

### Supplementary Figure S4

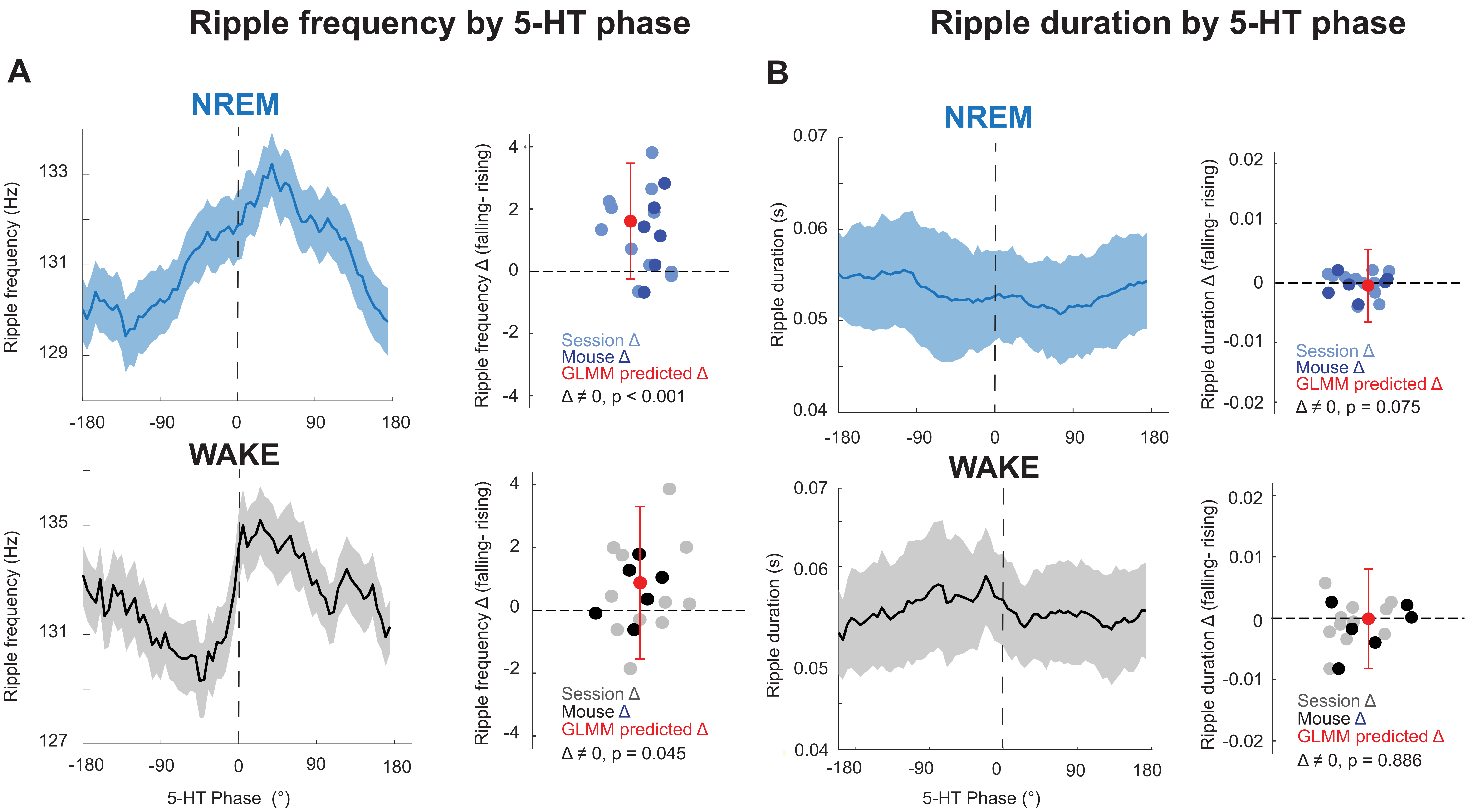
